# Phage-Plasmids Are Rare in Bacteria, May Exhibit Pseudolysogeny and Lack Antibiotic Resistance Genes

**DOI:** 10.1101/2025.11.07.687105

**Authors:** Gomathinayagam Sankaranarayanan, Kodiveri Muthukaliannan Gothandam

## Abstract

Phage-plasmids (PPs), hybrid mobile genetic elements possessing characteristics of both plasmids and bacteriophages, have recently gained attention for their proposed role in horizontal gene transfer, particularly in the dissemination of antimicrobial resistance genes (ARGs). In this study, we re-evaluated the prevalence, structure, and functional attributes of PPs across publicly available datasets comprising over 5 million viral and plasmid sequences. By employing a conservative and domain-centric approach that strictly filtered for high-quality phage genomes devoid of insertion sequences (ISs) and containing plasmid hallmark domains, we identified 3,002 putative PPs—representing a significantly lower proportion (∼0.25%) than previous estimates. Our functional analyses revealed that PPs are more closely related to virulent phages than to temperate phages or plasmids and primarily encode partitioning proteins rather than conjugative machinery, suggesting episomal maintenance and vertical inheritance. PPs displayed genome sizes larger than most phages or plasmids, indicating a potential fitness cost to their bacterial hosts and explaining their rarity. Despite prior claims, we found no evidence of ARG carriage in PPs or virulent phages; only a minority of temperate phages harbored such genes. Furthermore, the dihydrofolate reductase genes commonly mistaken as ARGs were excluded due to their structural and functional roles during phage infection. Interestingly, majority of the PPs were *Caudoviricetes* with plasmid partitioning proteins, and PPs classified under *Faserviricetes* universally carried the relaxase *NicK*, likely reflecting their rolling-circle replication rather than conjugative potential. Our findings challenge earlier generalizations regarding PPs and support a revised view that emphasizes their derivation from bacteriophages, limited functional resemblance to plasmids, and limited role in ARG dissemination.

## Introduction

In addition to resistance factor plasmids, phages have also been identified as vehicles for the transmission of antimicrobial resistance genes (ARGs) among bacteria. In 1961, Watanabe and Fukasawa documented the presence of phage p-22 in *Salmonella typhimurium* strain LT-2 and phage P1KC in E. coli strain K12, which were observed to transfer ARGs via phage conjugation(1). Initially referred to as ‘episomal resistance transfer factors,’ these findings underscored the role of phages in mediating ARG dissemination. Similarly, in 2010, Brenciani et al investigated bacteriophage Phim46.1, which harbored *mef*A and *tet*O genes conferring resistance to macrolides and tetracycline(2). Notably, these phages exhibited lytic activity, persisted as prophage in free circular form, and were capable of conjugally transferring ARGs. However, subsequent research suggested that phages infrequently encode ARGs which was previously reviewed by us in a prior article(3).

Traditionally, bacteriophages are characterized by their compact genomes. Occasionally, they may carry auxiliary metabolic genes, aimed at co-opting host machinery for their propagation rather than conferring mutual benefits to the host, but not ARGs(4, 5). Theoretically, carrying ARGs would increase the fitness chances of temperate phages inside their hosts. However, experiments evaluating bacterial uptake of prophages with and without ARG genes have found that bacterial hosts tend to favor either prophage or ARG acquisition, depending on environmental conditions (6). This phenomenon suggests that ARGs are selected against in phages. Bacteriophages possessing the capacity to integrate new genes may compromise their lytic cycle switching capability, leading to their transformation into cryptic prophages within host cells.

In contrast, plasmids are well-known carriers of ARGs and are often linked with antimicrobial resistance. Unlike phages, plasmids rely on host cells for replication and transfer but are capable of autonomous replication similar to phages. However, a distinguishing feature is their inability to exit host cells independently for horizontal transfer; instead, they necessitate contact with a compatible host cell to establish a conjugation apparatus for transfer(7).

Notably, there exist plasmids that lack autonomous transmission capabilities and instead rely on co-existing plasmids for conjugation apparatus and transmission, termed as non-conjugating or mobilizable plasmids. Indeed, the majority of plasmids fall into this category, with some carrying only a small region recognized by relaxase, known as *oriT* (origin of transfer)(8). Coluzzi et al. (2022) have reported that many plasmids lose their ability to conjugate and depend on the conjugation apparatus of co-existing plasmids. They also noted that conjugative plasmids may generate non-conjugative plasmids, which are often poorly adapted and prone to loss(7).

Consequently, non-conjugative plasmids are under pressure to encode accessory genes or be of particular size that confers a fitness advantage or indifference to the host, as otherwise, they may be eliminated due to fitness costs in subsequent host generations(7). Another intriguing aspect is the persistence of temperate bacteriophages within hosts, even when they confer no beneficial traits. To avoid fitness costs, these phages will often integrate themselves into the host genome.

At this juncture, a hybrid entity known as phage-plasmids has garnered attention among researchers. Although the term “phage-plasmids” was coined by Ravin et al. in 1999, it was Pfeifer et al. (year) who conducted extensive studies recently on the abundance of these entities in public databases and characterized them (9, 10).

Phages and plasmids typically exhibit distinct gene repertoires; however, when their gene sets overlap, they are identified as a distinct entity, namely phage-plasmids (PPs)(11). Additionally, the evolutionary origin of PPs is subject to debate: do they evolve from plasmids or phages? Initial reports by Pfeifer et al,(10) suggested a closer homology to phages, but subsequent evidence led to the revision that PPs are more akin to plasmids (12). In the intricate microenvironment within hosts, just to avoid incompatibility issues, plasmids can shed certain genes to become non-conjugative, on the other hand even minor alterations to phage genomes can render them non-functional. Thus, PPs must delicately balance their genetic makeup to persist in this transient state. A pertinent question arises regarding the presence of ARGs within this overlap. While Pfeifer et al’s earlier findings suggested that ARGs are more prevalent in PPs as compared to pure phages, we wanted to reassess this claim to understand what makes PPs special so as to carry more ARGs than the conventional phages(10, 13).

## Methods

### Collection of Databases

The genomes of phages and complete plasmids were sourced from various databases. Previously, Pfeifer et al. retrieved approximately 11,000 plasmids and 2500 complete phages from GenBank(10). However, we aimed to broaden our search to include other collections containing phage genomes assembled and retrieved from metagenomes. These databases include IMG-VR(14), ICTV bacteriophages(15), Refseq bacteriophages, Millard lab’s INPHARED 14April2025 collection of phages retrieved from GenBank (accessed on May 2025)(16), For negative controls, we retrieved phage genomes from *Cystoviridae* and *Leviviridae* from IMG-VR and BV-BRC(17). Complete plasmid sequences were retrieved from the plasmids database (PLSDB) (18).

### Collection of plasmid associated genes

Plasmid-associated HMM profiles were collected from various sources: 15 profiles from eggNOG(19), 56 profiles from TIGRFAM(20), 110 profiles from PFAM(21), 54 profiles from KEGG Orthology(22), 30 profiles from NCBI Clusters of Orthologous Groups, and 8 profiles from the Virus Orthologs database(23). These profiles were downloaded and used to create an HMM database. The profiles were collected based on information from tools such as MacSyFinder(24), ConjScan(25), and previously published articles(10, 13). Information on the HMMs from each of the respective databases can be obtained from Supplementary Table S1

### *oriT* Plasmid sequence search

Bacterial plasmid GenBank files were downloaded from NCBI Entrez. *oriT* sequences were extracted from the GenBank files based on the feature using an in-house Python script. The retrieved sequences and the experimentally verified nucleotide sequences from oriTDB were concatenated and used to create a BLAST database(8). The collected phage sequences were then searched for *oriT* sequences with a minimum percentage identity of 60% against the *oriT* BLAST database.

### Hallmark plasmid protein domain search in viral genomes

All computational analyses were conducted in a Unix-based environment using the Conda package manager to manage software dependencies. The tools employed included Prodigal-gv for Open Reading Frame (ORF) prediction(26), hmmsearch for protein homology searches against the custom HMM database(27), seqkit(28) and seqtk for sequence manipulation, and CheckV for quality control assessment of viruses(29).

Nucleotide sequences retrieved from all the above-mentioned databases were initially filtered to retain those exceeding 3,000 base pairs using seqkit. ORFs within these filtered sequences were then predicted using Prodigal-gv in metagenomic mode, generating protein sequences. Subsequently, these protein sequences underwent a homology search against the custom HMM database using hmmsearch, with an E-value threshold of 1e-20 to ensure robust matches. Corresponding nucleotide sequences for identified hits were retrieved from the original file via seqtk, underwent deduplication with seqkit, and quality control of these nucleotide sequences was conducted using CheckV’s ‘end_to_end’ mode, assessing completeness and quality. From the CheckV quality check, only sequences deemed high quality according to MIUViG (Minimum Information about an Uncultivated Virus Genome) standards and genomes with at least one viral hallmark gene were selected(30), while the rest were not taken for downstream analyses. The data was additionally filtered for HMM-lower bound based hits, excluding those and retaining other high and medium confidence viral gene annotations.

### Hallmark viral protein domain search in plasmids

Next, the retrieved plasmid sequences were filtered to retain only those with a length greater than or equal to 3,000 base pairs. Initially, phage-like plasmid sequences were predicted using the DeepVirFinder tool(31), and sequences with a score greater than 0.7 were selected for further analysis. The sequences filtered by DeepVirFinder were then subjected to CheckV analysis to assess completeness, following the same approach used for phage sequences. As with phages, the analysis excluded hits based on HMM lower-bound scores, retaining only those with high and medium-confidence viral gene annotations as identified by CheckV.

## Results

### Pfeifer et al’s Phage-Plasmids

To understand the characteristics of putative phage plasmids by Pfeifer et al., we downloaded the nucleotide sequences of these phage plasmids and searched for plasmid-hallmark proteins using a custom HMM database that we created. Out of the reported 780 phage plasmids, we identified hits for 681 phage plasmids from Pfeifer’s dataset. We then investigated the presence of insertion sequences within the putative phage plasmid hits, as it is unusual for bacteriophages to contain insertion sequences. Which resulted in 419 putative P-Ps without identifiable ISs in them. The same set of hits was subjected to quality analysis using CheckV. CheckV generates a set of result files in TSV format. It contains a contamination estimate file, which identifies whether the given sequences are prophages and determines loci of ‘islands’ based on the concentration of ‘phage’ or ‘host’ ORFs in the provided genomic sequences. We mapped the region coordinates of both insertion sequences and regions marked as host within the given genomic sequences and visualized the results and are presented in supplementary figure S1A,B.

From those images, it can be observed that some Pfeifer et al.’s phage-plasmid genomes are interspersed with several insertion sequences, and yet a few genomes do not have identifiable IS elements. Additionally, the ‘contaminant’ islands as identified by CheckV are highly localized. Typically, genomes with IS elements cannot be activated into phages. Therefore, it is pertinent to believe that these are no longer phages *sensu stricto*-which has lost the ability to activate into lytic phage and remains as cryptic prophage. Next, we analyzed how many of these reported PPs carry ARGs and *oriT* sequences. Only 47 of them carried ARGs, and very few of them contained the *oriT* sequence. These putative PPs with *oriT* sequences often carried other plasmid hallmark genes.

Another important observation is that there were also some PPs without identifiable IS elements and localized ‘contaminant’ ORFs. We wanted to investigate whether these could be PPs. To understand how typical PPs in the literature compare to these in these aspects, we conducted further analysis.

### Characteristics of Phage-Plasmids in earlier Reports

Next, we analyzed Ravin et al.’s popular phage-plasmid N15(9). As anticipated, bacteriophage N15 did not carry any IS elements, ORFs for integration and excision, or identifiable ARGs, but it did contain three moron/auxiliary genes. It is important to remember that N15 is a linear bacteriophage episomally found in its host.

Next, we analyzed Watanabe et al.’s genome contigs (E. coli K12 - GCA_028644645.1) in an identical manner. None of the contigs satisfied the set criteria for a high-quality phage, so they were excluded from further analysis(1).

Next in line were Piligrimova et al.’s ‘plasmid prophage sequences,’ which were examined for phage-plasmid characteristics based on the above criteria(32). Only one sequence (KP795655.1) was found to contain characteristics satisfactory for a PP (no IS elements, high-quality phage genome according to CheckV). Notably, this ‘plasmid prophage’ had no ORFs for integration and excision from the host genome, no ARGs, and no moron/auxiliary metabolic genes. Upon deeper gene-level analysis, we found that the approximately 43 kb ‘plasmid prophage’ carried proteins for plasmid partitioning (*rep*A and *Fts*K). Similarly, N15 also carried a gene for plasmid partitioning protein.

### Validation of Applied Methodology

To validate the correctness of the applied method, we applied it to the genomes of *Cystoviridae* and *Leviviridae* downloaded from NCBI, BV-BRC and IMG-VR, which were considered as negative controls. However, CheckV could identify only two complete genomes within *Cystoviridae*. Out of 3,091 sequences under *Leviviridae*, 817 were assessed as complete by CheckV. As anticipated, neither *Leviviridae* nor *Cystoviridae* contained IS sequences, plasmid hallmark proteins, or AMR genes. Based on the inference from all the above analyses, we proceeded to scrutinize the presence of phage-plasmids in sequences retrieved from databases, aiming to further characterize them by profiling their genomic and protein-level traits.

### Identification of Putative Phage-Plasmids

From a total of approximately 5.3 million sequences encompassing bacteriophages and plasmids, we identified 3,002 non-redundant putative PPs as shown in the table below. These PPs were defined as high-quality phage genome (MIUViG) - that contained at least one plasmid hallmark protein with a Pfam domain match at an E-value threshold below 1e-20.

We classified the identified PPs using the taxonomy identification module of Phabox. Out of 3,002 PPs, Phabox was able to classify 2,489. Among these, the majority were assigned to the class *Caudoviricetes* (82.3%), followed by *Faserviricetes* (16.5%) (Supplementary figure S2).

The same Phabox tool was used to predict host taxonomy. Most PPs were associated with bacterial hosts, primarily from the phyla *Proteobacteria* and *Firmicutes* (Supplementary Figure S3). Only a small number were predicted to infect archaeal hosts.

Regarding plasmid hallmark proteins present in the PPs, the partitioning proteins A, B, and C were the predominant domains in *Caudoviricetes*-classified PPs, with an average frequency of 0.32. In contrast, the average frequency of all other plasmid-associated domains in these PPs was 0.17. All PPs classified under *Faserviricetes* contained the DNA relaxase (*Nic*K) gene, while the frequency of other domains in this group was only 0.10. Similarly, all PPs assigned to *Megaviricetes* harbored the partitioning protein C (*parC*), and the mean frequency of other plasmid-associated domains in this class was 0.23.

In contrast, the occurrence of conjugation-related domains, such as those associated with the Type IV secretion system and the conjugal transfer proteins such as *tra*C and *tra*D, were sporadic among the identified putative PPs.

To aid in downstream functional comparisons, all other phage sequences (excluding the putative PPs) were classified into temperate and virulent categories using the Phabox tool(33), which predicts phage lifestyle based on gene content and regulatory elements. The phage dataset used for this classification was obtained from the earlier retrieved database (INPHARED) of Millard Lab.

Further, plasmid sequences that were not classified as putative PPs were categorized into three groups based on CheckV analysis: plasmids with multiple viral regions, plasmids lacking viral regions, and plasmids containing small viral islands. Plasmids were classified as lacking viral regions if no viral hallmark genes were identified. If a plasmid contained one or more viral hallmark region, it was designated as a plasmid with a small viral island. In cases where CheckV identified more than one viral genomic region within a plasmid based on k-mer profiling, the sequence was classified as a plasmid with multiple viral regions.

### Functional annotation and trait matrix construction

To investigate functional signatures across these six distinct groups, coding sequences (CDSs) were predicted again using Prodigal-GV and protein domains were detected using HMMER with a Pfam-A HMM profile and plasmid hallmark HMM profile previously used. Using a E-value threshold of 1e-5, domain annotations were binarized to generate a genome-by-domain trait matrix. This matrix served as the basis for further statistical analyses.

### Dimensionality Reduction and Clustering Reveal Functional Structure

To visualize functional similarity across genome groups, we performed Principal Component Analysis (PCA) on the trait matrix. PPs formed a semi-distinct cluster, partially overlapping with both virulent and temperate phages, while exhibiting minimal overlap with the plasmid categories (Figure 1A).

**Figure 1:**
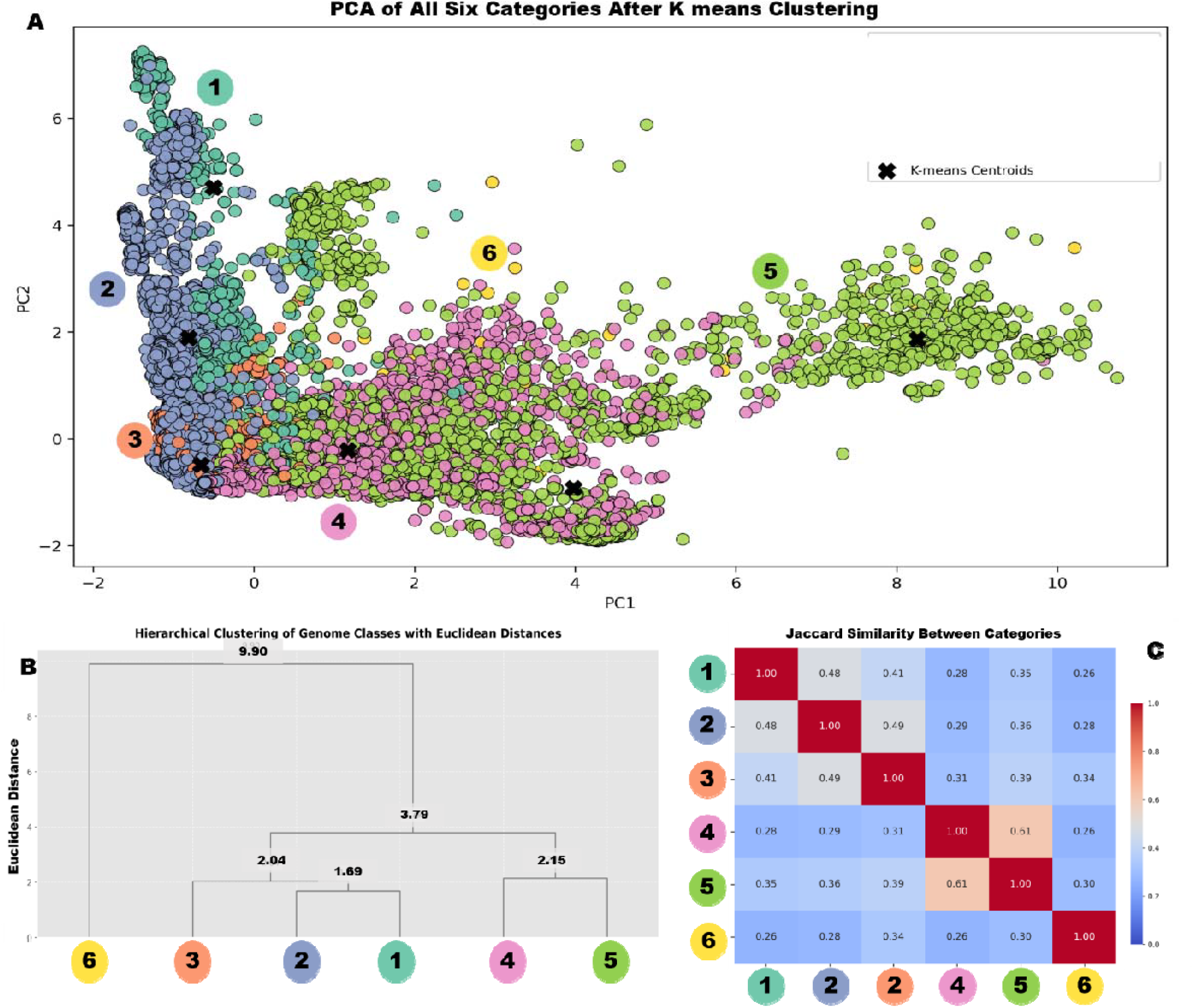
Functional comparison of genome categories based on protein domain profiles. 1) PPs, 2) Virulent phages, 3) Temperate phages, 4) plasmids lacking viral regions, 5) Plasmids with small viral regions, 6) Plasmids with multiple viral regions. (A) Principal Component Analysis (PCA) plot depicting the distribution of genome categories based on binarized protein domain presence-absence data. Phage-plasmids (PPs) form a semi-distinct cluster, partially overlapping with virulent and temperate phages, but showing minimal overlap with plasmid categories. (B) Hierarchical clustering dendrogram based on Ward’s method, constructed using the same trait matrix. PPs cluster more closely with virulent phages than with temperate phages or plasmids, while plasmids with multiple viral regions form a distinct clade. (C) Pairwise Jaccard similarity matrix illustrating domain-level overlap between genome categories. Higher similarity is observed between PPs and virulent phages, supporting their functional proximity

**Figure 2:**
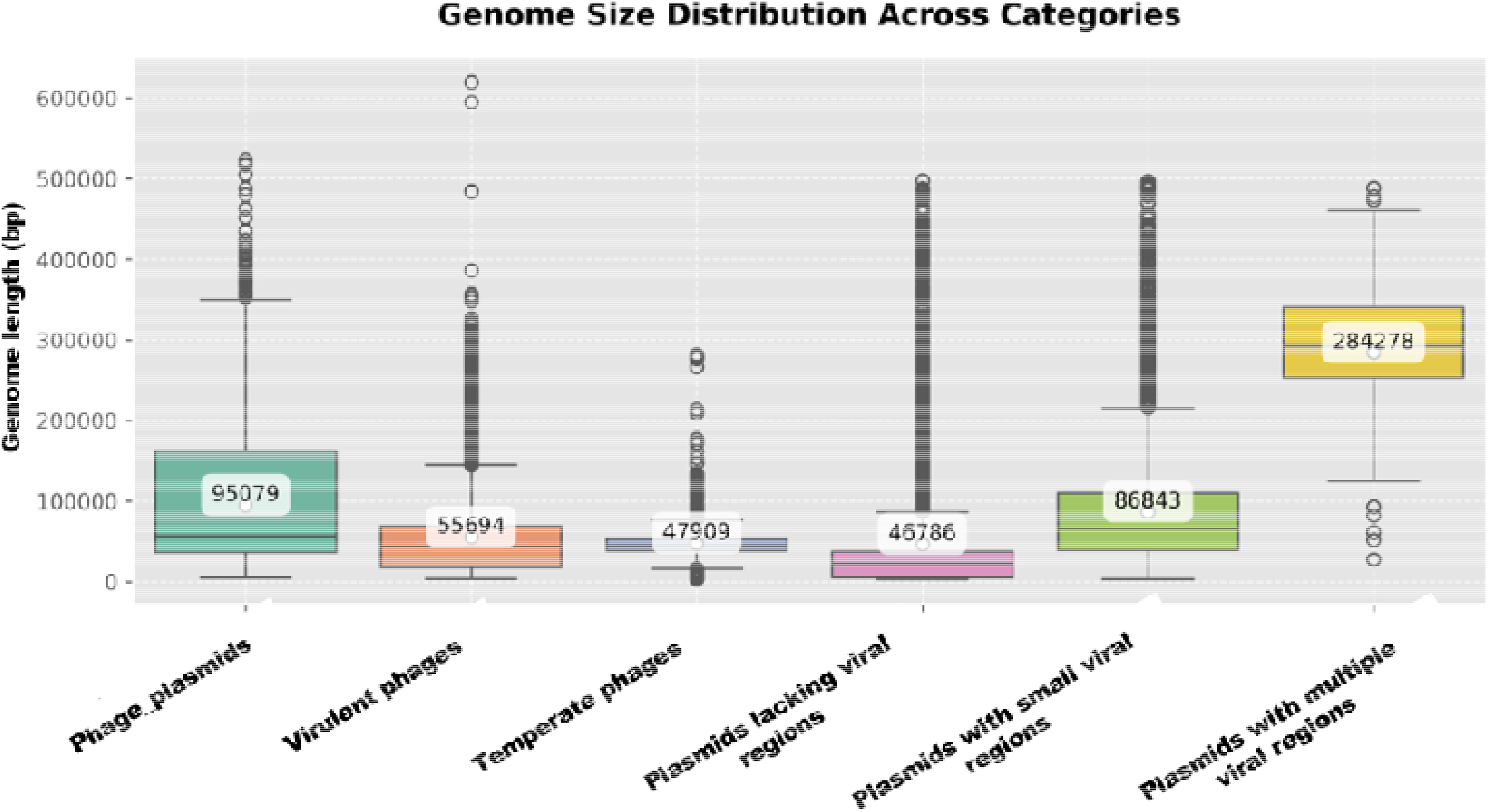
Genome size distribution across genome categories. Box-and-whisker plots showing the distribution of assembled genome lengths (in kilobases) across six categories: phage-plasmids, virulent phages, temperate phages, plasmids with no viral regions, plasmids with small viral regions, and plasmids with multiple viral regions. Plasmids with multiple viral regions exhibit the highest mean genome size (∼300 kb), while plasmids lacking viral regions have the smallest. Several phage-plasmids exceed the mean size of temperate and virulent phages.

**Figure 3:**
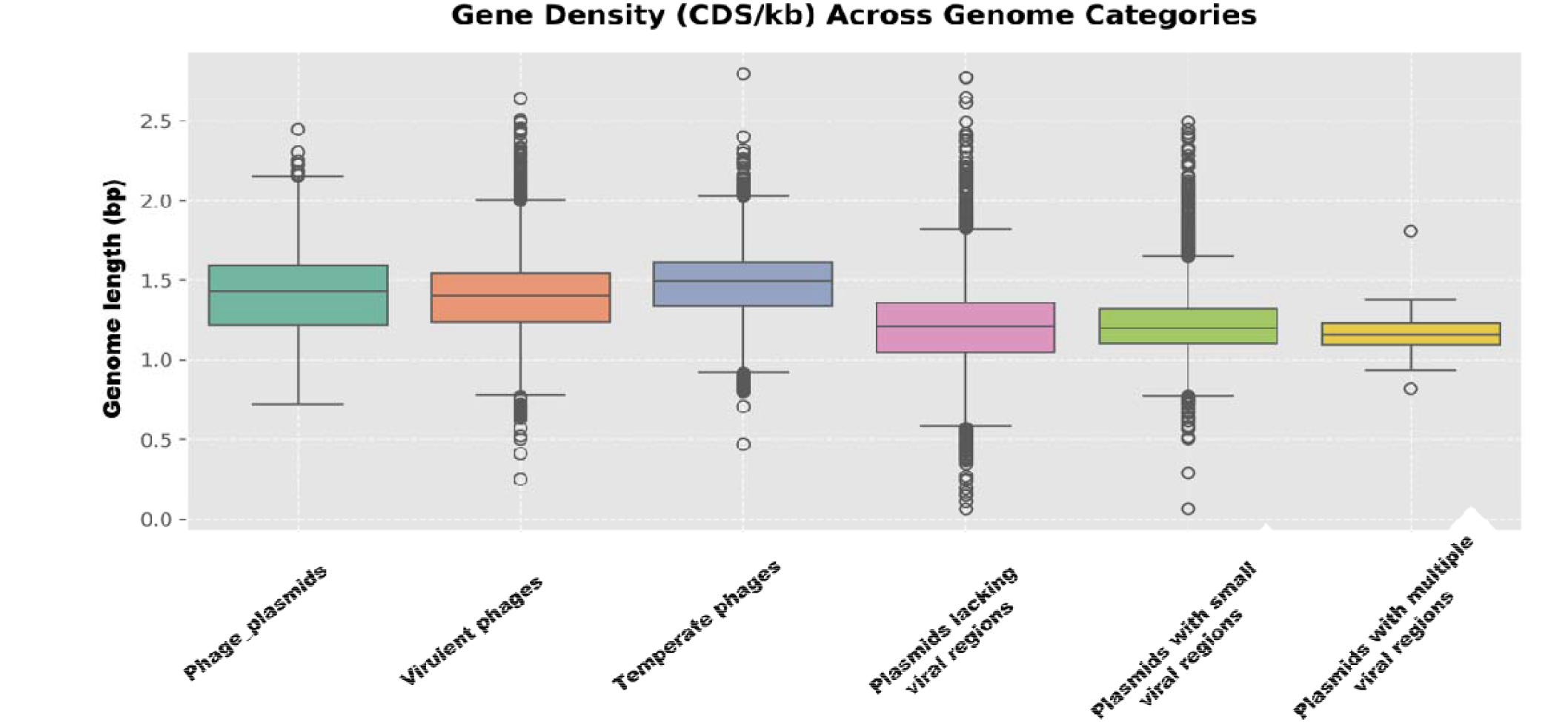
Gene density across genome categories. Box-and-whisker plots representing the number of coding sequences (CDSs) per kilobase of assembled sequence for each genome category. Phage-plasmids and virulent phages exhibit similar gene densities (∼1.4 CDSs/kb), while temperate phages show slightly higher density (∼1.5 CDSs/kb). All plasmid categories display lower gene densities (∼1.2 CDSs/kb)

To further examine the structure within the PCA space, we applied K-means clustering using the first two principal components. The number of clusters was set to match the number of predefined genome classes. K-means clustering explained 89% of the variance (variance explained = 0.89), indicating strong concordance between the inferred clusters and the underlying functional traits (Figure 1A). Notably, sequences classified as plasmids with small viral regions formed three distinct clusters, with their corresponding centroid clearly separated from the rest. Pairwise Jaccard similarities were then calculated between these category profiles, defined as the ratio of the intersection (shared domains) to the union (total unique domains) of domain sets (Figure 1C).

A Random Forest classifier trained on the trait matrix was used to identify Pfam domains that most effectively discriminated among the genome groups. The most predictive domains included phage integrases, plasmid partitioning proteins (e.g., K03496), plasmid replication initiation proteins, and conjugation-related domains.

Additionally, hierarchical clustering based on Ward’s method was performed using the same trait matrix(Figure 1B). The resulting dendrogram showed well-defined groupings, with PPs clustering more closely with virulent phages than with temperate phages or any of the plasmid categories. Another notable observation was that plasmids with multiple viral regions formed a distinctly separate clade, indicating substantial functional divergence from all other groups.

### Genome Size and Gene Density Analyses

To evaluate genomic characteristics across the defined categories, we analyzed both genome size and gene density based on predicted coding sequences (CDSs). Genome length was determined directly from assembled sequences in each category, while gene density was calculated as the number of CDSs per kilobase of sequence.

Among all categories, plasmids with multiple viral regions exhibited the highest mean genome size, averaging approximately 300 kilobases. In contrast, plasmids lacking any viral regions had the smallest mean genome size (Figure). The mean genome sizes of PPs, temperate phages, and virulent phages were broadly similar; however, several PPs displayed genome sizes that exceeded the mean values and were notably larger than the average temperate or virulent phage.

With respect to gene density across categories, PPs and virulent phages exhibited a mean gene density of approximately 1.4 CDSs per kilobase, while temperate phages had a slightly higher mean gene density of around 1.5. In contrast, all plasmid categories displayed lower gene densities, averaging around 1.2 CDSs per kilobase. For reference, the typical gene density of *Escherichia coli* is approximately 0.9 CDSs per kilobase.

### AMR gene profile

From the PPs, virulent phages, and temperate phages, we specifically excluded the trimethoprim class of antimicrobial resistance (AMR) genes conferred by dihydrofolate reductase (*dfr*) homologs. This exclusion was made because *dfr* genes in bacteriophages are dual-use proteins(34). During the lytic phase, the phage-encoded dihydrofolate reductase functions as a folate-reducing enzyme, redirecting host’s nucleotide synthesis toward viral genome replication. Subsequently, during phage assembly, the same protein becomes the tail baseplate structure and is released with phages upon host cell lysis.

A total of 95 *dfr* genes were identified in PPs, 18 in virulent phages, and 6 in temperate phages. These *dfr* genes, which confer resistance to trimethoprim, were excluded from downstream analyses due to their dual functional role in phage biology, as described previously. The above table presents the distribution of AMR genes excluding the trimethoprim class for PPs, virulent phages, and temperate phages. In contrast, the remaining three genome categories—plasmids with multiple viral regions, plasmids with small viral regions, and plasmids lacking viral regions include all AMR genes, including those of the trimethoprim class.

After excluding *dfr* genes, no antibiotic resistance genes were detected in PPs, and only one AMR gene was identified in virulent phages. The probability of detecting an AMR gene was highest in plasmids with multiple viral regions, followed by plasmids with small viral regions.

## Discussion

According to the classical modular theory of bacteriophage evolution, “each virus encountered in nature is a favorable combination of modules selected to work optimally both individually and collectively to fill a particular niche.”(35) Recent reports on PPs have extended this modular theory by introducing plasmids and bacteriophages as participants in this module exchange. These studies emphasize that PPs are hybrid in nature, which exhibit characteristics of both phages and plasmids. In line with that, several features have been proposed as defining traits of PPs: (i) PPs are intermediates between bacteriophages and plasmids; (ii) PPs facilitate gene flow between plasmids and phages, including the transfer of ARGs; (iii) PPs exchange genes more frequently with plasmids than with phages; (iv) some PPs lose genes over time and evolve into integrative prophages, whereas others acquire conjugation-related genes from plasmids and become mobilizable via conjugation; and (v) PPs tend to be larger than temperate phages(12).

Even within the framework of modular theory, although phage genomes exhibit mosaicism, not all genes within them are modular in nature. For instance, the head gene regions are generally not modular, unlike the tail regions, which are more frequently observed to be exchangeable(35). This constraint arises because the head genes encode proteins that interact intimately during the assembly of the phage capsid, and these proteins must co-evolve to maintain the structural and functional integrity of the head. Consequently, if recombination generates a hybrid genome by joining head gene modules from two different phages, the resulting recombinant is typically non-functional and thus eliminated from the population, despite each gene being functional in its original genomic context.

Moreover, phage genomes are characteristically terse and compact, with strong selective pressures maintaining this economy of sequence. Any disruption to this structural efficiency, such as insertions or the accumulation of non-essential sequences can compromise the phage’s viability. This is one of the reasons why transposable elements are counter-selected in bacteriophage genomes(5, 36). However, recent studies that have attempted to characterize PPs have largely overlooked this critical aspect of phage genome organization (10, 13). While it is true that some phage genomes may carry IS, this cannot be considered a general or defining feature of functional phages. The presence of such elements in a genome mostly imply that the phage is inactive or incapable of entering the lytic cycle.

In this study, this principle has been applied *sensu stricto*, whereby PPs containing IS were categorically excluded from consideration. Furthermore, we did not employ any machine learning techniques to classify PPs from publicly available databases. Instead, classification was based on the presence of plasmid-hallmark protein domains within phage genomes. As for differentiating PPs from plasmids, we relied on the CheckV tool, which was utilized to identify hallmark viral genes within plasmid sequences. CheckV is capable of demarcating mosaic genomic regions of viral or host (in this context, plasmid) origin within a given sequence, thereby enabling the identification of prophage regions within bacterial genomes.

This approach has resulted in a substantial reduction in the number of PPs identified from publicly available databases, contradicting the earlier estimate by Pfeifer et al., who reported that PPs constitute approximately 5% of total phages and 7% of plasmids. In contrast, our analysis indicates that only 0.25% of entries in public databases excluding metagenome-derived phages from IMG-VR qualify as *bona fide* PPs. Notably, no PPs could be identified within the PLSDB plasmid sequence database. While several plasmids were found to contain ‘mosaics’ of viral regions, these are better considered as degenerate phages now being plasmids, as they are unlikely to represent viable phages.

This distinction represents a significant confounding factor in earlier studies that characterized PPs as intermediates between bacteriophages and plasmids and proposed that PPs exchange genes more frequently with plasmids than with phages. A key finding of our study, based on domain trait analysis, is that PPs show stronger alignment with virulent phages than with temperate phages or plasmids. Another claim made in prior studies was that PPs may lose genes to become integrative phages and simultaneously acquire conjugation-related genes to adopt plasmid-like mobility. However, it is important to recall that plasmid types follow a source–sink model, where only approximately 25% of plasmids are conjugative. These conjugative plasmids often shed their conjugation machinery, giving rise to mobilizable plasmids that may retain only *oriT* sequences and a relaxase.

Given this context, it is unlikely that PPs would readily acquire conjugation modules from plasmids. This is further supported by our observation that conjugation-related domains such as those associated with the type IV secretion system were only sporadically detected in the identified PPs. To test whether this pattern might be an artefact of the stringent HMMER threshold (E-value ≤ 1e−20), we relaxed the threshold to 1e−5. However, this adjustment did not result in a meaningful increase in the detection of conjugation-associated domains (data not shown). Another noteworthy observation is that PPs, in order to remain viable phages, would likely need to acquire genes enabling their integration into host genomes not vice versa.

However, one observation is consistent with earlier findings that the average genome size of putative PPs identified in our study was larger than that of most bacteriophages and plasmids. Only plasmids harboring multiple viral regions exhibited a higher mean genome size. The size of an episome is a well-established factor influencing the fitness cost imposed on the host; larger episomes tend to impose a greater metabolic burden and are therefore subject to counter-selection. This constraint may partially explain why PPs are so infrequently observed among bacterial genomes.

Additionally, the majority of the identified putative PPs were classified as *Caudoviricetes*, despite the inclusion of phage sequences derived from metagenomic sources. These *Caudoviricetes* PPs were strongly associated with plasmid partitioning proteins rather than conjugative components, suggesting a preference for episomal maintenance and vertical transmission rather than horizontal gene transfer. Although they share similarities with virulent phages, these PPs likely exhibit a form of temperate behavior, allowing entry into daughter cells without inducing host cell lysis.

Interestingly, the second most prevalent taxonomic group among the identified PPs was *Faserviricetes*. All *Faserviricetes* PPs encoded *nicK*, a gene for DNA relaxase. In plasmids, relaxase is typically associated with the presence of an *oriT* sequence. However, our BLAST-based analysis identified only a single *oriT* sequence, and it was not found in a *Faserviricetes* genome. It is important to note that the currently available *oriT* sequence datasets are not comprehensive. A plausible explanation for the ubiquitous presence of the relaxase gene in *Faserviricetes* PPs is that *Faserviricetes* are single-stranded DNA (ssDNA) phages. Upon host entry, they convert to a double-stranded form and replicate via a rolling-circle mechanism, akin to plasmid replication, and are released from the host by budding rather than lysis. In this context, relaxase plays an essential role in the replication process during the pseudolysogenic phase and is not related to conjugation.

Another important aspect of this study is to assess the reported association between ARGs and PPs. Previous studies identified episomes exhibiting phage-like characteristics that harbored ARGs(10, 13). Notably, Pfeifer et al. estimated that PPs contained more ARGs than phages but fewer than plasmids. This observation may be attributable to two key factors. First, Pfeifer et al. may have included false-positive PPs in their ARG assessments. Second, it is critical to recognise that commonly used annotation tools such as AMRFinder(37) and CARD-RGI classify dihydrofolate reductase (*dfr*) genes as ARGs.

However, in the context of bacteriophages, *dfr* genes often serve dual roles: they function as structural proteins and play a crucial role in manipulating the host’s nucleotide synthesis during the lytic cycle. Many studies have attributed the presence of *dfr* to trimethoprim resistance, leading to generalized claims that bacteriophages contribute to the dissemination of ARGs. Such interpretations may overlook the biological context in which these genes function, thereby overstating the role of phages particularly PPs in ARG propagation.

In our investigation, we found that AMR genes were entirely absent from both PPs and virulent phages, and were identified in only ∼2% of temperate phages. In contrast, plasmid categories, particularly those containing viral proteins showed a markedly higher probability of harboring AMR genes, as demonstrated in Table 2. This observation underscores that the presence of viral signatures within plasmids is more strongly associated with AMR gene carriage than is the case for bona fide plasmids without detectable viral hallmark genes.

**Table 1:**
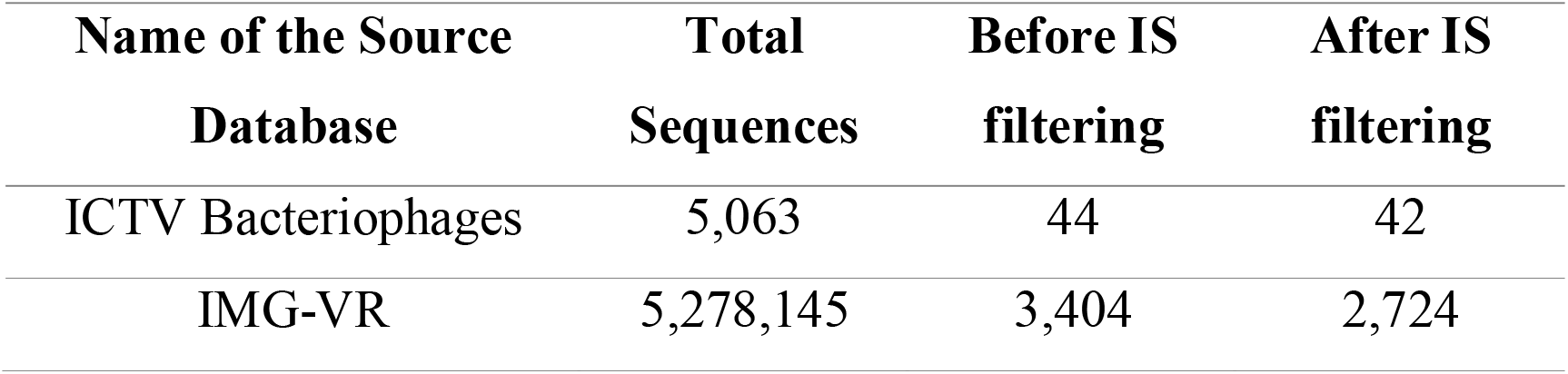
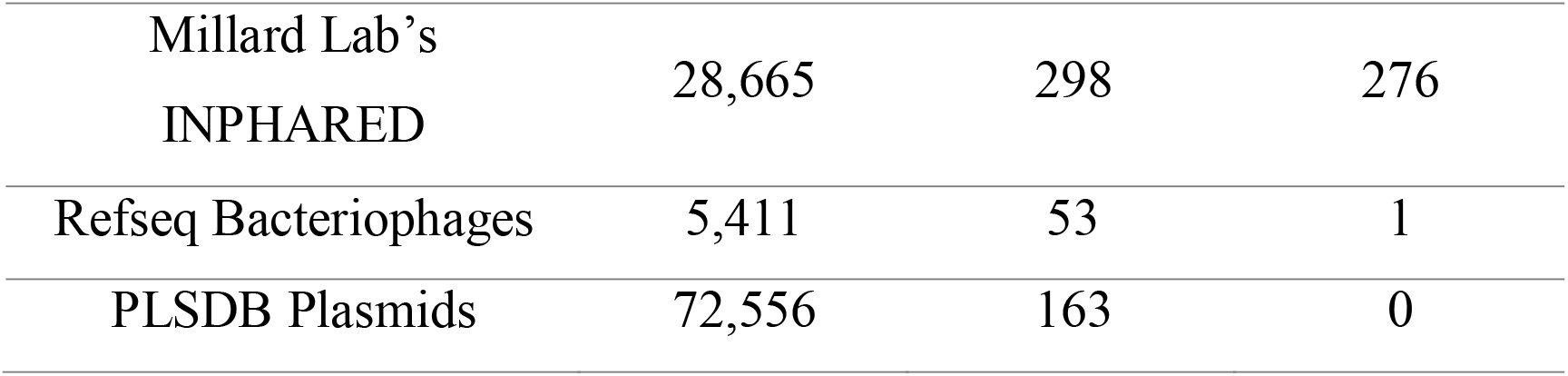
Summary of Sequence Counts from Source Databases Before and After Insertion Sequence Filtering.

**Table 2:** Distribution of Antimicrobial Resistance (AMR) Genes Across Different Categories.

| Category | Total Genomes | Genomes with $\geq 1$ AMR | Total AMR Genes | AMR Gene Load per Genome | AMR Gene Presence Probability |
| --- | --- | --- | --- | --- | --- |
| Phage_plasmids | 3002 | 0 | 0 | 0.0000 | 0.0000 |
| Virulent_phages | 19304 | 1 | 1 | 0.0001 | 0.0001 |
| Temperate_phages | 6198 | 121 | 134 | 0.0216 | 0.0195 |
| Plasmids_with_mvr<sup>1</sup> | 112 | 89 | 490 | 4.3750 | 0.7946 |
| Plasmids_lvr<sup>2</sup> | 5683 | 2080 | 4160 | 0.7321 | 0.3660 |
| Plasmids_with_svr<sup>3</sup> | 6270 | 4100 | 10537 | 1.6805 | 0.6541 |

1-Plasmids with multiple viral regions, 2 – Plasmids lacking viral regions, 3- Plasmids with small viral regions.

## Conclusion

Based on our analysis, PPs are more closely related to virulent phages than to temperate phages and are clearly distinct from plasmids. They are more frequently found as episomes, and only a minority appear to exhibit conjugative properties. These elements may exhibit pseudolysogeny, either by delaying lysis until favorable conditions arise or by releasing progeny through budding rather than host lysis. Importantly, PPs do not show evidence of antimicrobial resistance dissemination, a characteristic shared with *bona fide* bacteriophages. While PPs may have acquired certain modular elements from plasmids, their evolutionary origin appears to be rooted in bacteriophages. In parallel, numerous plasmids have incorporated cryptic phage-derived modules, forming mosaic structures that should not be mistaken for active phage-plasmids or phages.

## Data availability statement

The basic scripts used for the analysis, along with all the HMM profiles, are available in the following GitHub repository: https://github.com/GomathiNayagam/Phage_plasmid_survey.

## Acknowledgement

We thank Dr Allison Elaine Mann from the Department of Anthropology, University of Wyoming for revising the initial draft and providing the computational resources used.

