## Supplementary figure for "Phage-Plasmids Are Rare in Bacteria, May Exhibit Pseudolysogeny and Lack Antibiotic Resistance Genes"

Supplementary Figures

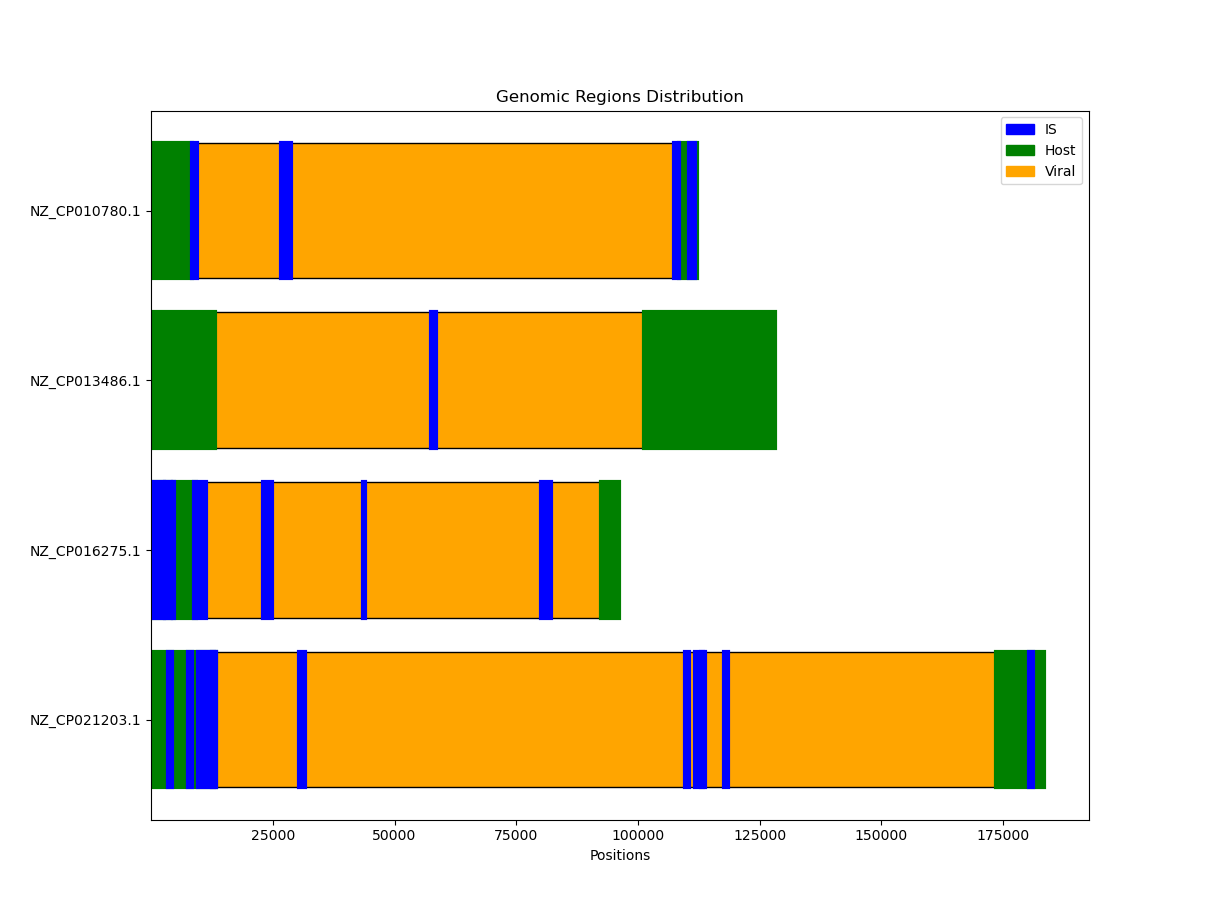

Figure S1A

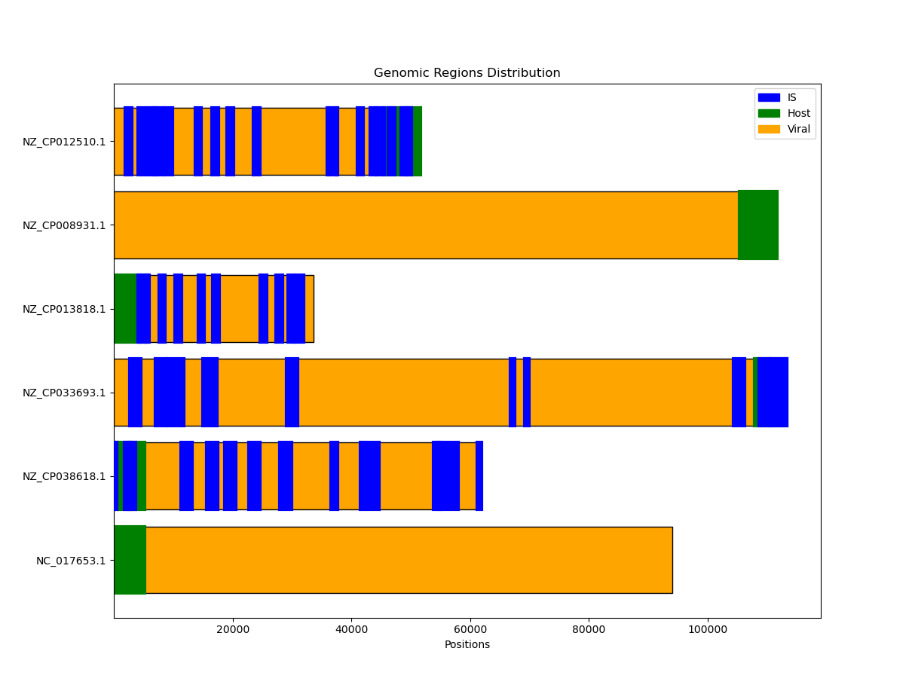

Figure S1B Linear genome maps of Pfeifer *et al’s* phage-plasmids showing ‘host’ modules and several interspersed insertion sequences indication transposable elements.

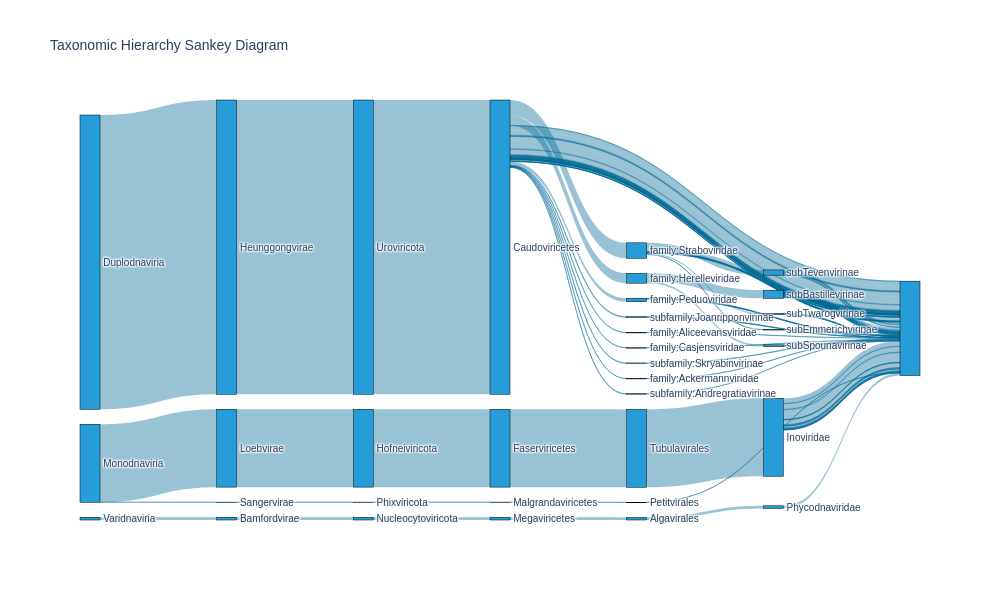

Figure S2. Sankey diagram of taxonomic classification of phage-plasmids using Phabox

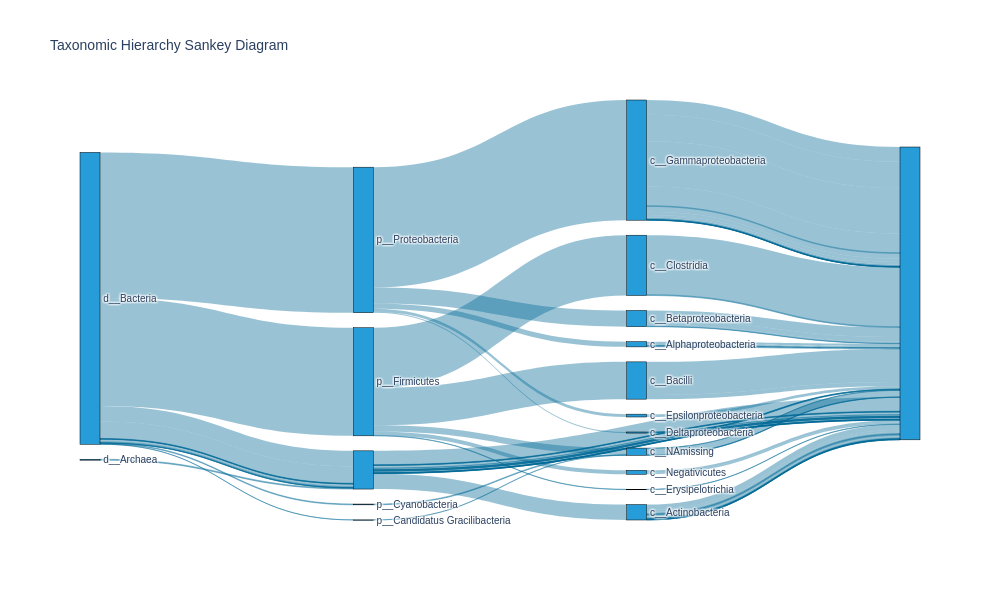

Figure S3. Sankey diagram of taxonomic classification of predicted hosts of phage-plasmids using Phabox.
